# Unethical amnesia brain: Memory and metacognitive distortion induced by dishonesty

**DOI:** 10.1101/2024.03.03.583239

**Authors:** Xinyi Julia Xu, Dean Mobbs, Haiyan Wu

## Abstract

Unethical actions and decisions may distort human memory in two aspects: memory accuracy and metacognition. However, the neural and computational mechanisms underlying the metacognition distortion caused by repeated dishonesty remain largely unknown. Here, we performed two fMRI studies, including one replication study, with an information-sending task in the scanner. The main moral decision task in the scanner involves consistency and reward as two main factors, combined with a pre-scan and post-scan memory test together with mouse tracking. With multiple dimensions of metrics to measure metacognition, we test whether the inter-subject metacognition change correlates with how participants trade off consistency and reward. We find that the compression of representational geometry of reward in the orbitofrontal cortex (OFC) is correlated with both immediate and delayed metacognition changes. Also, the functional connectivity between the dorsolateral prefrontal cortex (DLPFC) and the left temporoparietal junction (lTPJ) under dishonest responses can predict both immediate and delayed metacognition changes in memory. These results suggest that decision-making, emotion, and memory-related brain regions together play a key role in metacognition change after immoral action, shedding light on the neural mechanism of the complex interplay between moral decisions, cognitive processes, and memory distortion.

*‘I did that’, says my memory*.

*‘I could not have done that’, says my pride, and remains inexorable.*

*Eventually - the memory yields*.

—Nietzsche[^1^]

## Introduction

Immoral behaviors, like dishonesty, may exert powerful influences on both performance and feeling of memory, as they can result in psychologically uncomfortable feelings such as guilt and shame^1^. Because of the threat to self-perception^2^, people may exhibit unethical amnesia - a self-protective mechanism by reducing the dissonance and stress of socially unacceptable behavior^3^. Such dynamic processes of memory retention and retrieval can be monitored by metacognition^4–6^. Metacognition about memory refers to one’s belief in their abilities or knowledge regarding a specific memory content^7–9^, which may be unrelated to objective performance^10^. One approach to measuring metacognition is harnessing statistical signatures of self-reported confidence in a self-centered frame of reference^9, 11, 12^. A prominent signature for this measurement is the folded-X pattern, which indicates that confidence should increase with signal strength for correct trials, and decrease with signal strength for error trials. People with a high degree of metacognitive sensitivity would show more confidence when they are right, and less confidence when they are wrong^13^. This pattern is consistent in both human and animal^11, 14^, and has been used as a marker of confidence-related physiological and neural signals^15^.

Studies on morality and memory have revealed that people forget the instances where they lied or made selfish choices^3^. Moreover, laboratory lie detection studies offer evidence showing that guilty participants can suppress retrieval of their behaviors, and generate biases of eyewitness memory^16^ or autobiographical memories^17^. One leading interpretations stem from cognitive dissonance theory^18^, which suggests that discrepancies between truth and conflicted responses lead to psychological discomfort. Indeed, substantial evidence suggests that forgetting is one way of cognitive dissonance reduction^19^. For example, Elkin and Leippe (1986) discovered that dissonance-related arousal, as assessed physiologically, did not diminish after an attitude change. However, when participants have time to forget about the dissonance, a decline in its level is observed^20^. The results raise an interesting question about whether repeated immoral behaviors lead to forgetting, with false memories that can not be discriminated against.

A prominent perspective of repeated immoral behaviors is that people adapt to it (e.g., the aversive affects of immoral behavior weaken with repeated acts^21, 22^). From this perspective, repeated dishonesty is like a general emotional memory, which could be two-fold. First, the immoral event usually accompanies negative feelings and is typically more vivid^23, 24^. On the other hand, people who act unethically are more likely to suppress their memory to protect their self-image, which results in decreased memory accuracy or vividness^25^. This view is supported by a recent study that unethical events would become more blurred or rated as less morally wrong^26^. However, another effect, coined as the illusory truth effect^27^, indicates people’s tendency to believe false information to be correct after repeated exposure^28, 29^. Although such truth illusion exists, its metacognitive experiences (e.g., confidence) after unethical decisions and the neural mechanism are still less understood. One potential approach is incorporating mouse tracking (MT) to capture metacognition or confidence^30^. Mouse tracking provides a continuous response mode, incorporating richer information than binary response mode^31^. It has also been applied to measure memory strength and confidence^32^ or detect memory with autobiographical implicit association test (aIAT)^17^. Accordingly, mouse tracking can capture the post-effect driven by dishonesty, which could be conceptualized as both the memory changes in accuracy or confidence and the hesitation during the decision.

To manipulate moral conflict setting, reward and consistency, are two critical components in dishonesty^33^. Most motivated dishonesty studies are attributable to reward or self-interest^34^, and neuroimaging studies imply the involvement of the reward brain region in dishonesty^35–39^. For instance, self-serving dishonesty accompanies increased striatum activity and stronger functional connectivity (FC) between valuation (e.g., orbitofrontal cortex, OFC) and cognitive control brain networks^37^. It should be noted that the dopaminergic memory consolidation hypothesis supports that extrinsic rewards may facilitate long-term memory^40^, which may interplay with the moral decision itself. During this dynamic process, the hippocampus may play a key role as it has been shown that memory items are represented via item-specific distinct patterns of brain activity during the encoding period and dynamically reactivated, or distorted during the recall period^41–43^. On the other hand, when faced with repeated situations, the desire to be and appear consistent is a powerful determinant of human behavior, which can be viewed as a type of keeping self-image consistent^44^. However, rare research examines the effect of self-consistency and reward on moral decisions and the neural association of their post-effect on memory(immediate vs. days later).

In the present study, with a complementary manipulation of pre-task and post-task measures, and with three conditions in the moral decision(e.g., *enhance dishonesty, enhance honesty, and random*)(see Section **Study design and procedure in Figure 1A**), we investigate (i) whether moral decisions lead to accuracy and metacognition change, and (ii) if so, how this change is associated with cognitive control and memory neural system. We predict the influence of immoral decisions on memory metacognition on memory would be evident by decoupling respective contributions of reward and consistency to behavior and its neural basis. Specifically, we have the following predictions: (i) following the dishonest responses, items that participants being dishonest in the information passing task would show a decline in overall memory performance and metacognition when they tested the memory immediately or days after; (ii) the trade-off process and brain activity in dishonesty decisions are associated with memory distortion via distorted representation processing, which is independent of the visual or semantic features. To identify brain activity related to metacognition change of memory, we utilized multiple-step tasks with both out-scanner memory tests and an information-sending task (IST task in the fMRI scanner. Our focus is on the metacognition changes, but also consider the brain activity during decision and memory performance after the behavior decisions. Through various metrics of assessing metacognition, we find that metacognition in memory is altered following immoral responses. This metacognition change is predicted by reward compression of OFC in neural geometry, during moral decisions.

**Figure 1.**
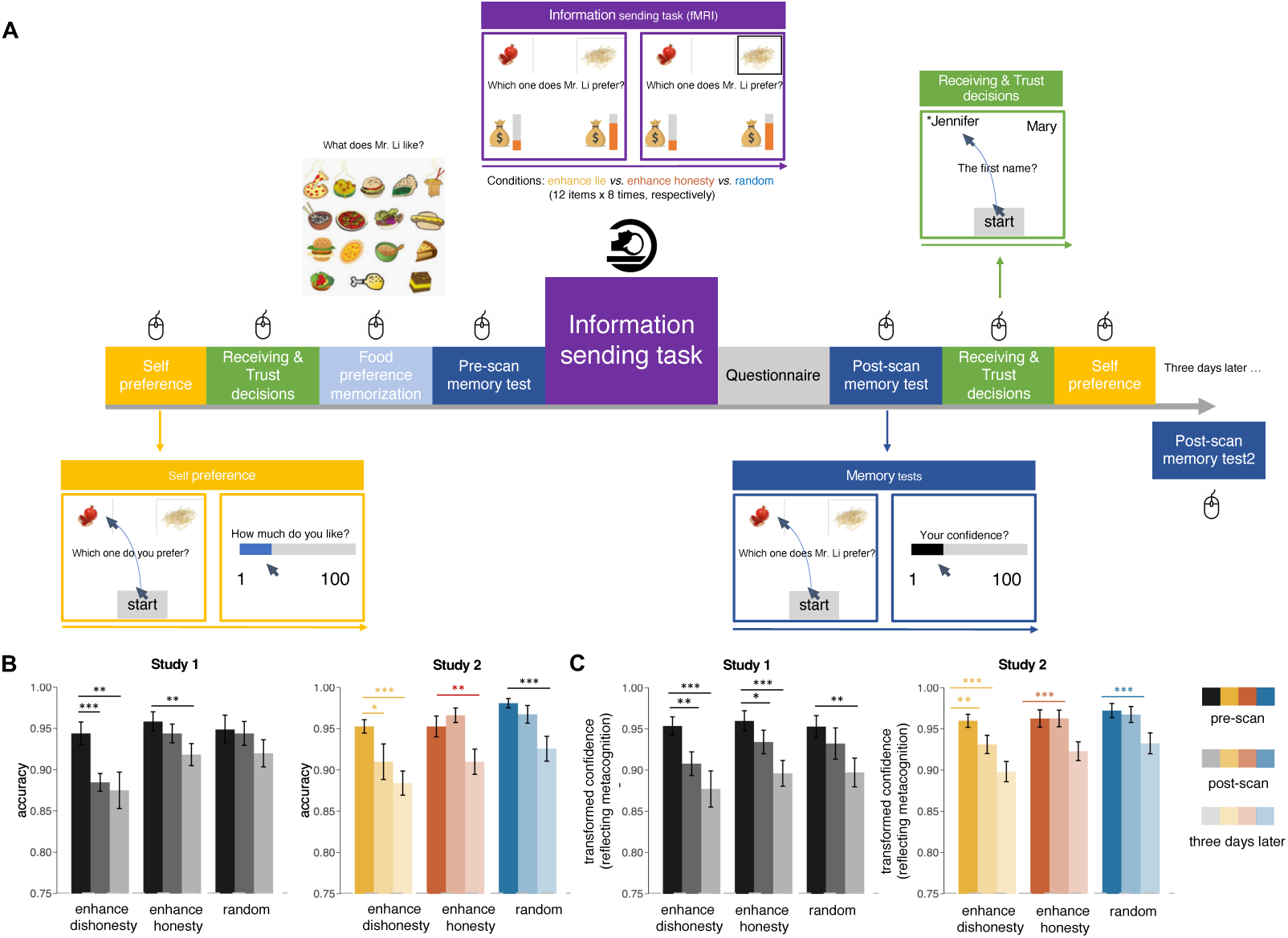
The experimental design and behavioral performance. **(A)** For each participant, there were 9 parts (8 parts for day 1 and 1 part for three days later) for the whole study. Participants performed a food memory task and a food preference task before and after the Information Sending Task (IST). IST was conducted in the fMRI scanner, where participants were asked to pass the preference information of Mr. Li to the next participant (the receiver) with the consideration of reward units in four scan sessions. We manipulated reward unit difference (referred to as “reinforcer”in this paper) between two items and kept them lying about specific items (repeated dishonesty) or making truthful responses with 4 sessions of fMRI scanning. **(B)** The accuracy and **(C)** transformed confidence (using the quadratic scoring rule to quantify metacognition level, see *Methods*) in pre-scan, post-scan and three-day-later memory tasks. The transparency of color represents the period of memory tests.

## Results

### Study 1

#### Memory accuracy and transformed confidence declined after repeated dishonesty

Participants were arranged to perform a 9-part task (Figure 1A) which consisted of two self-preference tests, two trust tests, and three memory tests separated by the information-sending task (IST, conducted in fMRI scanner). We first described the memory accuracy and self-reported confidence change before and after the IST. Because the accuracy of pre-scan memory test needed to reach 90% accuracy to start the IST, both the accuracy and confidence score matched across conditions (*enhance dishonesty* condition, *enhance honesty* condition and *random* condition) in this part (accuracy: *F*_2,75_ = 0.245, *p* = 0.783; 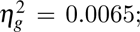 confidence: *F*_2,75_ = 0.099, *p* = 0.91; 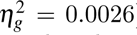). After the IST, participants performed the 2nd and 3rd memory tests right after the scanning and three days later. Interestingly, the accuracy of memory only declined significantly under the *enhance dishonesty* condition in the first post-session test (*enhance dishonesty*: *t* = 4.59, *df* = 25, *p <* 0.001***, 95% CI from 0.033 to 0.086; Figure 1B left panel; for statistical results, see Table S5 in supplementary materials). From the first pre-scan test to the three-day-later test, there was a significant memory accuracy decline under both *enhance honesty* and *enhance dishonesty* conditions (Figure 1B left panel; Table S5 in supplementary materials).

From pre-scan to post-scan, transformed confidence reflecting metacognition ability (for detailed calculation, see next section and *Methods*) changed under both *enhance honesty* and *enhance dishonesty* conditions, while from the first pre-scan test to the three-days-later test, confidence decline occurred under all conditions (Figure 1C, left panel; for statistical results, see Table S7 in supplementary materials). On the group level, we examined whether confidence could track accuracy, and found only in the pre-scan memory task participants’ confidence ratings were correlated with accuracies (Figure S7A). When we examined the self-reported confidence, significant changes occurred under both *enhance honesty* and *enhance dishonesty* conditions (*enhance honesty*: *t* = 1.97, *df* = 25, *p* = 0.06, 95% CI from -0.16 to 7.74; e*nhance dishonesty*: *t* = 1.97, *df* = 25, *p* = 0.06, 95% CI from -0.19 to 8.21). From the first pre-scan test to the three-day-later test, confidence decline occurred under all conditions (Figure S4A, left panel; for statistical results, see Table S9 in supplementary materials).

To examine whether transformed confidence change was related to response randomness, we used entropy^45^ as a measure of the overall randomness of participants’ choices under different conditions. On the subject level, we filtered items with no time-out trial. On the item level, four types of transition probability for responses were conducted to calculate the choice entropy (for details, see supplemental materials). Then, we averaged the choice entropy across items as an assessment of the randomness^33^. First, the results showed that there was a significant main effect of condition on choice entropy (*F*_2,75_ = 34.35, *p <* 0.001; 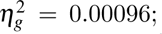 Figure S4C). To examine the relationship between transformed confidence change and choice randomness, we performed a linear mixed effect model with randomness as a fixed effect and subject as a random effect for *enhance dishonesty* and *random* condition separately. We found no relationship between transformed confidence change and response randomness under both conditions (*enhance dishonesty* & pre-post: *beta* = 0.013, *p* = 0.43; *random* & pre-post: *beta* = 0.018, *p* = 0.21; *enhance dishonesty* & pre-three days later: *beta* = -0.020, *p* = 0.37; *random* & pre-three days later: *beta* = 0.036, *p* = 0.07).

For **trial-by-trial ‘consistency’** (which was referred to across the article), we defined it as the difference of cumulative history responses between incorrect choice and correct choice. We defined ‘reward’ as the difference of reward between incorrect choice and correct choice. Three conditions of reward were designed to manipulate the value of consistency, that is, cumulative history response. However, the *current reward* only affected *future consistency* in the next run, so current reward and current consistency could be viewed as two independent variables. Previous study^33^ showed that both consistency and reward affected the dishonesty responses, we plotted the dishonesty probability as the function of reward and consistency, which was displayed in Figure S4D. This suggested that the participants considered both reward and consistency when they chose to lie or not in the IST.

#### Multidimensional measurements reflected metacognition change

In addition, multidimensional measurements be constructed as an index of metacognition change, as the self-reported confidence changes of some participants were slight (see Figure S6 in supplementary materials). To get more insights into implicit metacognition, we calculated the mouse tracking measurements, e.g., MAD, AD, and AUC, for the assessment of metacognition. The reaction time (RT) was also taken into consideration because it has been proven to be correlated with confidence^46^. Because metacognition describes the ability of how closely the confidence tracks performance, according to the folded-X pattern, we transformed the variables above using the quadratic scoring rule (QSR) to jointly measure the level of metacognition^11, 12, 47, 48^(see *Methods*). Significant decline of transformed variables from pre-scan to post-scan and three-days-later test were observed in the *enhance dishonesty* condition (see Figure S5 in supplemental materials), while the changes of untransformed variables were unstable (see Figure S3 in supplemental materials). Also, to validate whether transformed RT and mouse tracking measurements reflected metacognition, we performed a linear mixed effect model which showed that RT and mouse tracking measurements could predict transformed confidence in every stage of the memory test, and their changes could also predict transformed confidence change (Figure S9 left panel in supplemental materials). Then, changes from pre-scan to post-scan and from pre-scan to the three-days-later test of such five transformed indices were averaged among trials per subject and used to measure metacognition change together (Figure 4A). Because of the co-linearity between mouse trajectory measurements and the different scales between five transformed indices, we conducted principal component analysis (PCA) on these indices and selected the first three components to evaluate metacognition change (which was referred to across the article) as they explained over 90% of variance (Figure 4A). As expected, measurements from confidence ratings, RTs and mouse tracking indices all reflected metacognition after transformation.

#### Cognitive control brain was involved in the Information Sending Task (IST)

To understand the neural processes involved in the conflicting interplay between consistency and reward in repetitive moral decision-making, we sought to dissociate the two processes from the fMRI data of IST. Previous studies^49, 50^ suggest that cognitive control is necessary to override the dominant honest impulses to cheat for interest. Thus our primary focus was on cognitive control-related and reward-related brain regions of interest (ROIs). To determine whether our IST activated specific brain regions, we simply estimated a whole-brain GLM under dishonesty and honesty conditions. Contrast to honesty condition, PrACC, vlPFC, DLPFC and retrosplenial cingulate cortex (RSC1) were tracked under the dishonesty condition (RSC1: β = -6.75 ± 13.46, *t* = -2.51, *p* = 0.019; prACC: β = 1.55 ± 3.51, *t* = 2.24, *p* = 0.034; vlPFC: β = 1.97 ± 4.62, *t* = 2.17, *p* = 0.040; DLPFC: β = 2.37 ± 4.54, *t* = 2.66, *p* = 0.013; Figure 2A upper panel). In whole-brain analyses, three significant regions (DLPFC, preSMA, and insula) were found for dishonesty vs. honesty responses and PCC was activated in honesty VS. dishonesty responses (all *p_uncorrected_ <* 0.01; Figure 2B; for FDR corrected results, see Table S11 in supplemental materials). Moreover, for trials where the reward probability of incorrect choice was higher than correct choice, we didn’t observe a significant difference in reward-related ROIs like OFC (β = 2.64 ± 6.84, *t* = 1.97, *p* = 0.060; Figure 2C upper panel), but in cognitive control ROIs (MTG: β = 2.73 ± 4.93, *t* = 2.82, *p* = 0.009; DLPFC: β = -4.72 ± 4.26, *t* = -5.65, *p <*; lTPJ: β = 3.04 ± 7.47, *t* = 2.07, *p* = 0.049; see Figure 2C upper panel).

**Figure 2.**
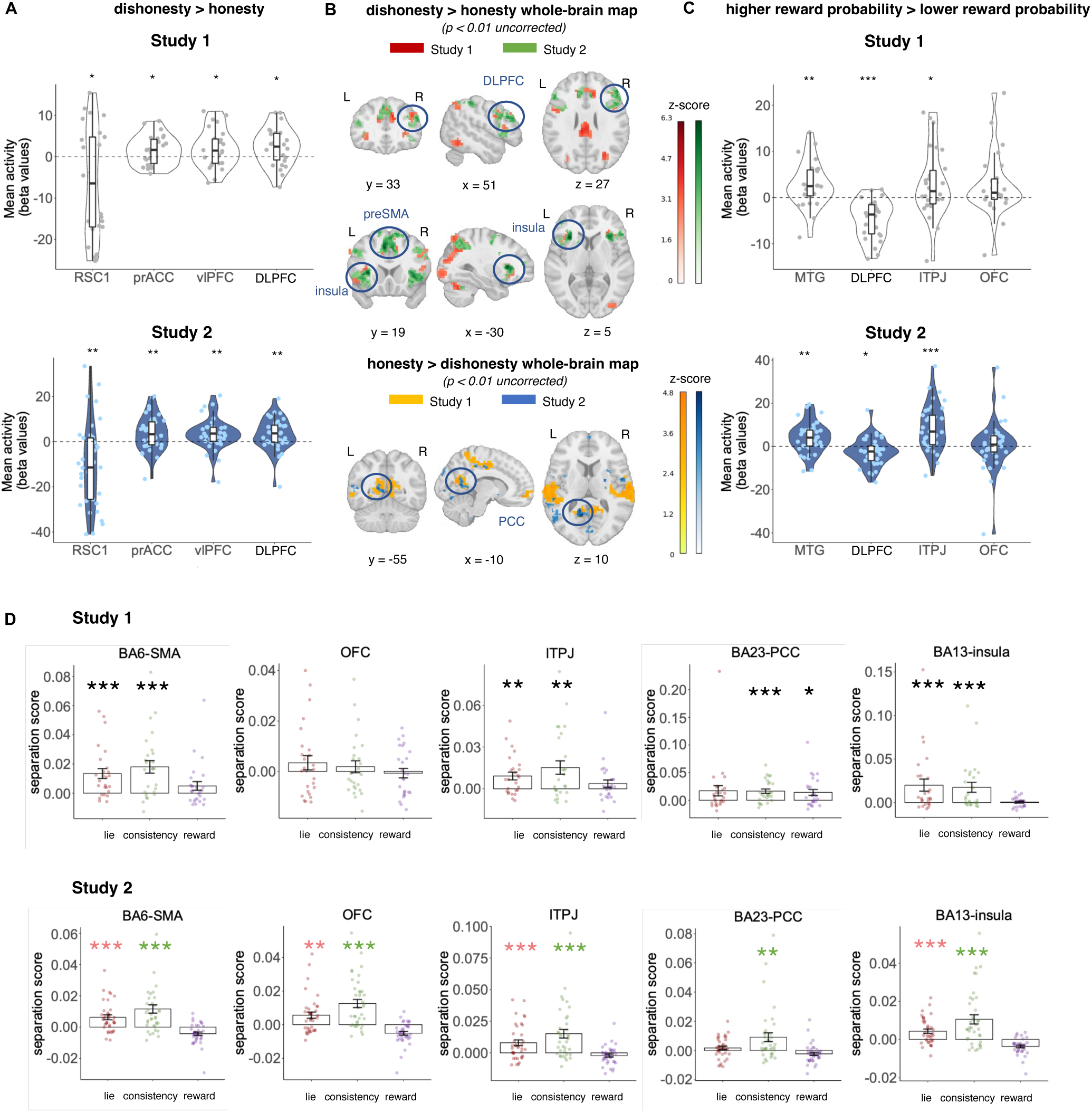
The univariate fMRI analysis results in study 1 and study 2. **(A)** Replicable mean beta values of ROIs under dishonesty condition compared to honesty condition, indicating greater activity over brain regions associated with cognitive control for dishonesty responses **(B)** The whole-brain beta map under dishonesty condition compared to honesty condition revealed stronger brain activity over regions related to cognitive control, like DLPFC and PCC across two studies. **(C)** Mean beta values of ROIs under the condition where the reward probability of incorrect choice was higher than correct choice. We didn’t find the activation of OFC for reward in both studies. **(D)** Separation score (between-condition similarity minus within-condition similarity) of dishonesty, consistency, and reward within the ROIs. The separation effect for consistency was observed in lTPJ, PCC, and insula, indicating cumulative dishonesty responses were represented in memory and cognitive control systems.

#### Cumulative dishonesty responses were represented in memory and cognitive control systems

We first examined the activity elicited by consistent dishonesty in consideration of the responses (dishonesty vs. honesty) and the explicit manipulation of reward probability. **Separation scores** were calculated as the average dissimilarity in activity elicited across conditions (consistency *>* 0 vs. consistency *<* 0) compared to the average dissimilarity within the condition. Hence, a positive separation score indicates that an area encodes representations of consistency. The separation effect was significant in both cognitive control areas like DLPFC (*df* = 25, *t* = 2.29, *p* = 0.031) and dACC (*df* = 25, *t* = 3.34, *p* = 0.0013; Figure S11D middle panel), emotion systems like lTPJ (*df* = 25, *t* = 3.15, *p* = 0.002) and insula, and memory system like hippocampus (*df* = 25, *t* = 2.44, *p* = 0.01; Figure S11D right panel in supplemental materials).

Separation scores for dishonesty and reward probability were calculated and analyzed across conditions and time stages. However, we didn’t observe the separation effect for dishonesty in OFC (*df* = 25, *t* = 1.23, *p* = 0.116; Figure 2D upper panel) and PCC (*df* = 25 *t* = 1.87 *p* = 0.0738; Figure 2D upper panel). Further, the separation effect for reward was observed in DLPFC (*df* = 25, *t* = 1.72, *p* = 0.0485; see Figure S11 in supplementary materials) and PCC (*df* = 25, *t* = 2.71, *p* = 0.01; Figure 2D), indicating most ROIs exhibited similar multi-voxel patterns of gaining reward versus not gaining reward.

#### Compressional rotated neural geometry of consistency and reward under dishonesty and dishonesty responses

If consistency and reward were represented differently under dishonesty and honesty responses in the brain, then this should be apparent if we compare the representations for different conditions. All the trials were split into several conditions according to the combination of the value of consistency differences (-7 to 7; might be different for each participant), reward differences (dishonesty as 1 and honesty as 0), and responses (dishonesty as 1 and honesty as 0). We put these three factors into a 3-dimensional space. Adapted from Flesch et al.^51^, we constructed seven representation model RDMs, which yielded 12 model RDMs in our case (see *Methods* in supplemental materials for details). For each participant, we constructed the neural RDMs among these conditions from the GLM-3 beta maps (see *Methods* for details). We found that in lTPJ, the second orthogonal model could explain the neural RDMs (*t* = 2.93, *p* = 0.043; see Figure 3A). However, in SMA, OFC and insula, more than one model explained a significant amount of variance in the neural RDMs, while in PCC no model could explain the neural RDM (Figure 3A; see Table S15 in supplementary materials).

**Figure 3.**
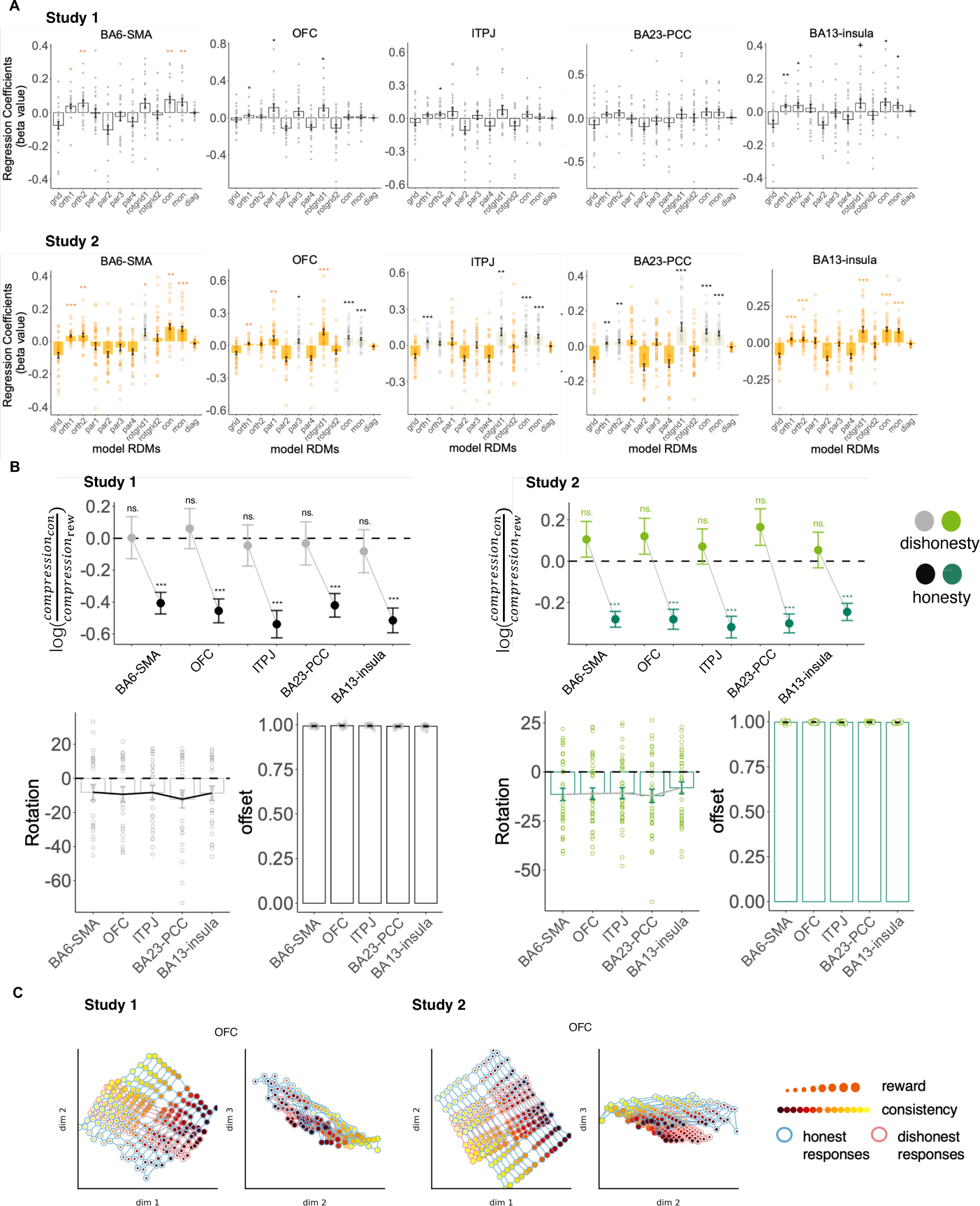
Results of neural geometry analysis of the IST fMRI data. All the trials were split into several conditions according to the combination of the value of consistency differences (-7 to 7; might be different for each participant), reward differences (dishonesty as 1 and honesty as 0), and responses (dishonesty as 1 and honesty as 0). We put these three factors into a 3-dimensional space. **(A)** We constructed seven representation model RDMs, which yielded 12 model RDMs in our case (see main text and *Methods* in supplemental materials for details). For each participant, we constructed the neural RDMs among these conditions from the GLM-3 beta maps (see *Methods* for details). Twelve model RDMs were fitted to five independently defined ROIs, showing response-agnostic encoding. In lTPJ, the second orthogonal model could explain the neural RDMs (*t* = 2.93, *p* = 0.043). However, in SMA, OFC and insula, more than one model explained a significant amount of variance in the neural RDMs, while in PCC no model could explain the neural RDM. **(B)** We built RDMs by varying these factors continuously, rather than fitting models encoding extremes of compression, rotation, and context separation. We yielded six parameters: 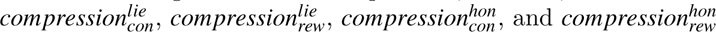 controlled for the compression along consistency and reward dimension under dishonesty and honesty responses respectively; the *rotation* parameter controlled for the response-dependent rotation of the variable axes (consistency and reward) from native space into the reference frame of the response; the context *o f f set* parameter controlled for the parallel distance between honesty and dishonesty responses. The fitting results showed compression along reward was significantly larger under dishonesty in all ROIs (SMA: *df* = 25, *t* = *−*6.01, *p <* 0.001; OFC: *df* = 25, *t* = *−*6.08, *p <* 0.001; lTPJ: *df* = 25, *t* = *−*6.28, *p <* 0.001; PCC: *df* = 25, *t* = *−*5.69, *p <* 0.001; insula: *df* = 25, *t* = *−*6.63, *p <* 0.001). **(C)** Low-dimensional projections of fMRI data taken from OFC, reconstructed from coefficients of the parameterized model described in **(B)**. The color of circles in the grids ranging from black to yellow corresponds to the value of reward difference ranging from -8 to 8, and the size of circles from small to large represents the value of consistency difference rangning from -7 to 7.

Thus, we built RDMs by varying these factors continuously, rather than fitting models encoding extremes of compression, rotation, and context separation. We yielded six parameters: 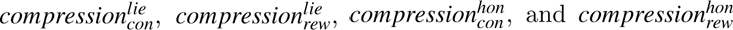 controlled for the compression along consistency and reward dimension under dishonesty and honesty responses respectively; the *rotation* parameter controlled for the response-dependent rotation of the variable axes (consistency and reward) from native space into the reference frame of the response; the context *o f f set* parameter controlled for the parallel distance between honesty and dishonesty responses. Fitting this parameterized model (see *eq.*5 in *Methods*) to the neural data and conducting t-tests for the log ratio of compression parameter for consistency and reward confirmed that compression along reward was significantly larger under dishonesty in all ROIs (SMA: *df* = 25, *t* = *−*6.01, *p <* 0.001; OFC: *df* = 25, *t* = *−*6.08, *p <* 0.001; lTPJ: *df* = 25, *t* = *−*6.28, *p <* 0.001; PCC: *df* = 25, *t* = *−*5.69, *p <* 0.001; insula: *df* = 25, *t* = *−*6.63, *p <* 0.001; Figure 3B upper left panel). For honesty responses, the compression of consistency and reward didn’t show a significant difference (Figure 3B upper left panel; see Table S17 in supplementary materials for details). The estimated rotation parameter was not close to zero (SMA: *M* = -8.17; OFC: *M* = *−*9.37; lTPJ: *M* = *−*8.33; PCC: *M* = -12.18; insula: *M* = -8.67; Figure 3B left lower panel), which suggests that the information of consistency and reward was not kept in the frame of reference of the inputs, yielding un-orthogonal representations in the ROIs. This ought to be that, because the cumulative history responses were due to the manipulation of reward in each session. The third dimension encoded responses, indicated by a non-zero offset parameter (SMA: *M* = 0.99; OFC: *M* = 0.99; lTPJ: *M* = 0.99; PCC: *M* = 0.99; insula: *M* = 0.99; Figure 3B left lower panel). Taking OFC as an example, the final neural geometry was visualized by multidimensional scaling (MDS) (Figure 3C left panel), where the length of reward in spaced grids of dishonesty condition was shorter than consistency.

#### The compression of reward under dishonesty was associated with metacognition change

To examine whether such compression of reward in the brain would lead to metacognition change, we conducted Spearman’s rank-order correlations between inter-subject metacognition change and the compression ratio of dishonesty responses. It showed significant correlations between compression ratio under dishonesty responses and metacognition change of pre-scan to post-scan test in SMA, OFC, lTPJ, PCC and insula (SMA: *rho* = 0.10, *p* = 0.045; OFC: *rho* = 0.16, *p* = 0.0025; lTPJ: *rho* = 0.18, *p <* 0.001; PCC: *rho* = 0.16, *p* = 0.0027; insula: *rho* = 0.14, *p* = 0.0073; Figure 4B upper panel). For metacognition change from pre-scan to three-days-later test, only OFC was found significantly correlated with the compression ratios (OFC: *rho* = 0.27, *p <* 0.001; Figure 4C upper panel).

**Figure 4.**
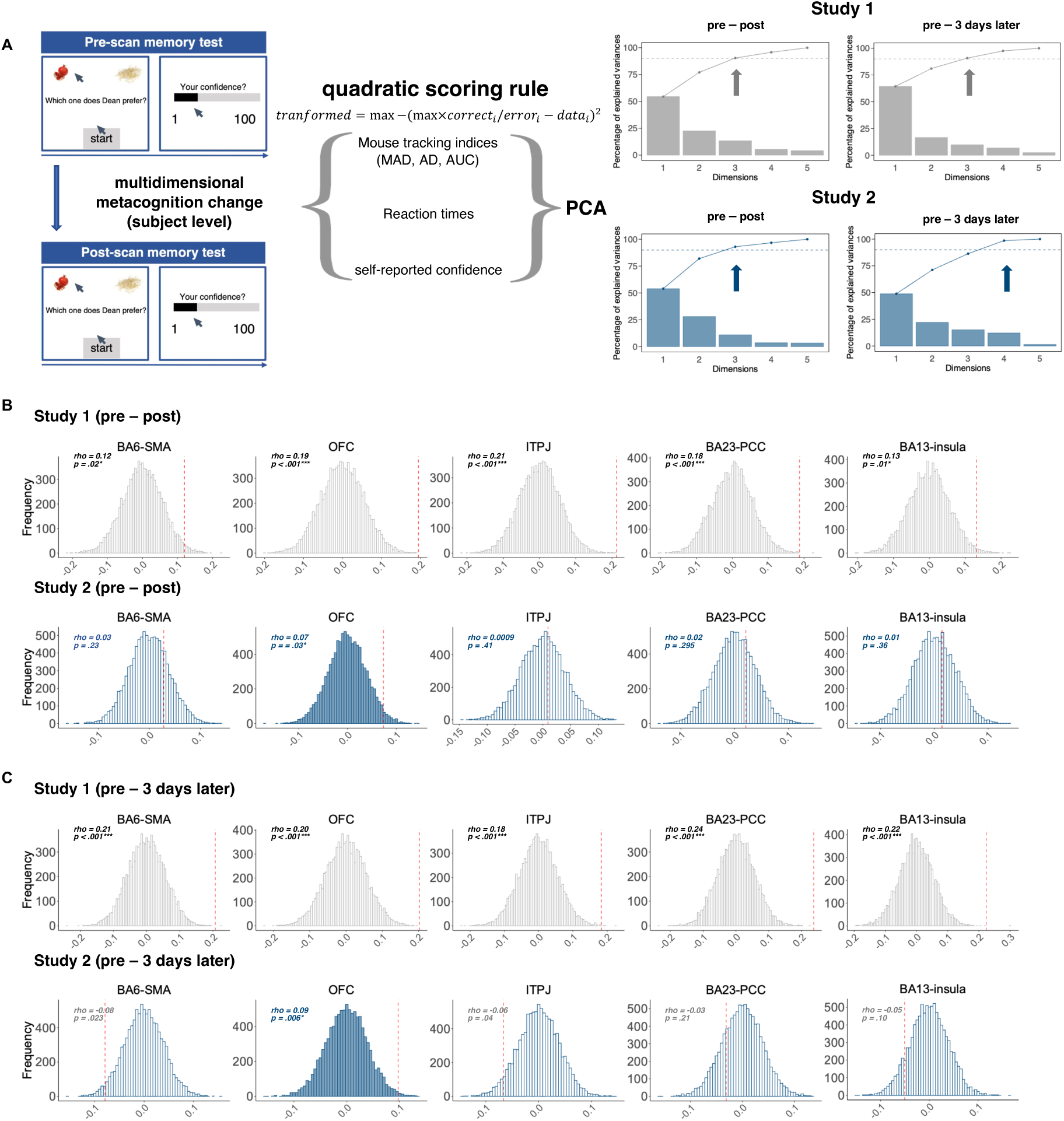
**(A)** Self-report confidence ratings, mouse tracking indices and reaction times were transformed using the quadratic scoring rule to quantify the multidimensional overall metacognition. The change of each transformed measurement from pre-scan test to post-scan test and three-day-later test was calculated on subject level. Considering the collinearity among these measurements, we conducted principal component analysis (PCA). Then the first three/four components were selected to evaluate metacognition change as they explained over %90 of variance. **(B)** Results of inter-subject representational similarity analysis. Permutation results of Spearman’s correlation between immediate multidimensional metacognition change and compression ratio under dishonesty responses. Only compression in OFC was correlated with both immediate and **(C)** delayed metacognition change.

#### Functional connectivity between DLPFC and lTPJ was associated with metacognition change

Playing a key role in cognitive control and dishonesty^52^, DLPFC was selected as a hub in functional connectivity(FC) analysis. Results showed significant FCs between DLPFC and PCC, insula and hippocampus (Figure 5A, left panel), echoing the involvement of DLPFC in memory change. Further, we examined whether there were task-related FCs between DLPFC and other ROIs that could predict the degree of metacognition change. We used inter-subject RDMs and implemented linear regression to explore the extent to which FCs between DLPFC and other ROIs could predict metacognition change variation (Figure 5C). The results showed that the FCs between DLPFC and lTPJ significantly predicted immediate and delayed metacognition change (immediate: β = 1.31, *p <* 0.001, 95% CI from 0.69 to 1.92; delayed: β = 0.69, *p* = 0.03, 95% CI from 0.078 to 1.29; Figure 5B).

**Figure 5.**
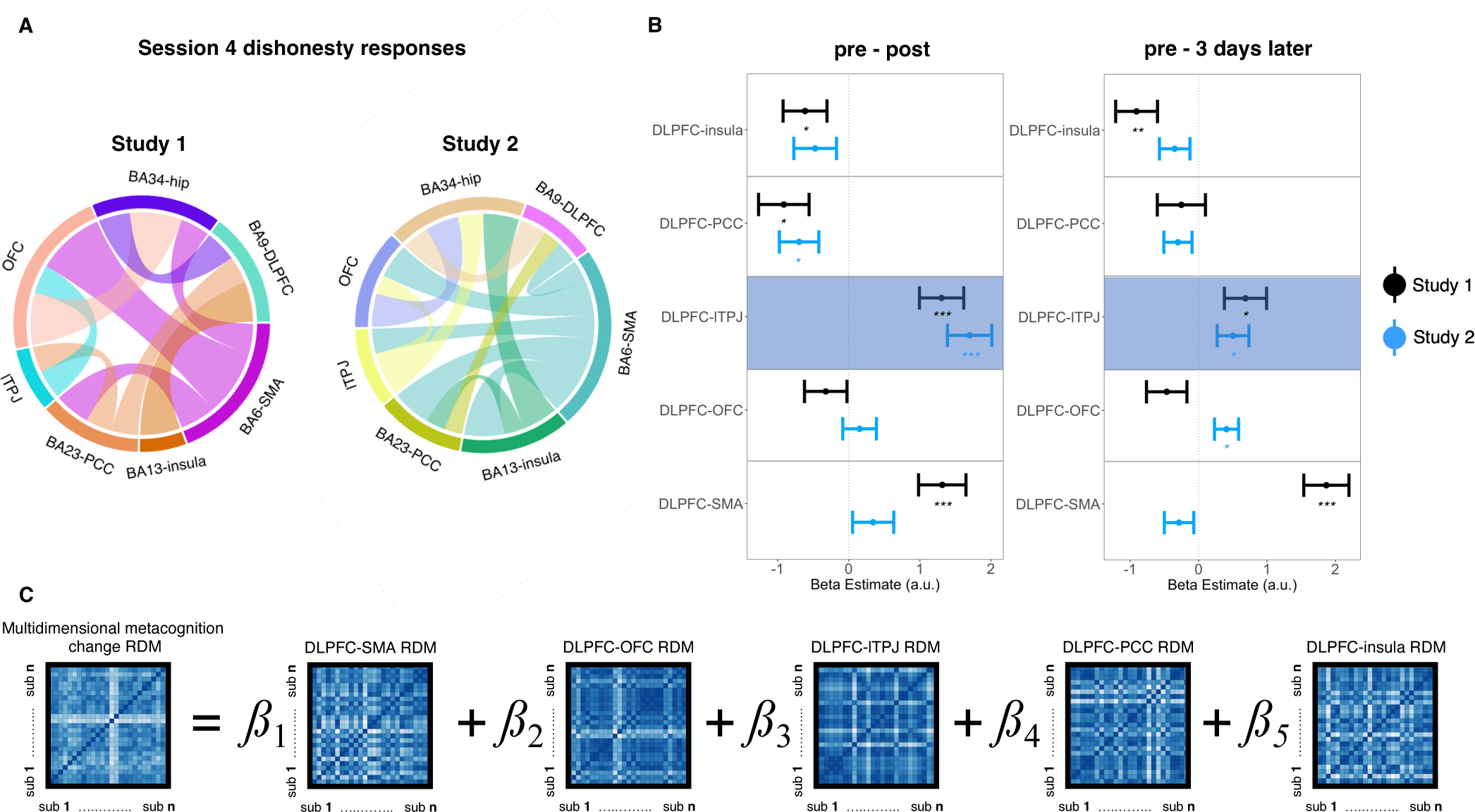
The functional connectivity analysis and results. **(A)** For both study 1 and study 2, the functional connectivity between DLPFC and PCC, hippocampus were significant. **(B)** The coefficients of the regression model shown in **(C)**. The coefficients of functional connectivity between DLPFC and lTPJ were significant when predicting later immediate and delayed metacognition change, and could be replicated across two studies. This indicated that metacognition change induced by repeated dishonesty extended beyond local, retrospective monitoring. In all the plots, the data points reflected beta estimates and error bars reflect standard error (* indicates *p <* 0.05, ** indicates *p <* 0.01, *** indicates *p <* 0.001). **(C)** The regression model to predict multidimensional metacognition change RDM.

### Study 2

#### Replication of memory change after the Information Sending Task (IST)

Similar as study 1, both the accuracy and self-reported confidence of memory test did not vary across conditions before the IST(*enhance dishonesty* condition, *enhance honesty* condition and *random* condition) in this part (accuracy: *F*_2,108_ = 3.075, *p* = 0.0503; 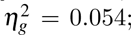 confidence: *F*_2,108_ = 0.15, *p* = 0.864; 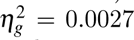). After IST, the accuracy of memory declined significantly under the *enhance dishonesty* condition in the post-scan test (*enhance dishonesty*: *t* = 2.18, *df* =36, *p* = 0.036*, 95% CI from 0.0029 to 0.083; *enhance honesty*: *t* = -1.78, *df* = 36, *p* = 0.08, 95% CI from -0.029 to 0.0019; *random*: *t* = 1.43, *df* = 36, *p* = 0.16, 95% CI from -0.0056 to 0.03; Figure 1B right panel). Also, the changes in transformed confidence which reflected metacognition ability under both the *enhance dishonesty* and *random* conditions were replicated (Figure 1C, right panel; for statistical results, see Table S8 in supplementary materials). On the group level, the results were replicated only in pre-scan memory task that participants’ confidence ratings could track accuracy after moving one outlier (Figure S7B). However, different from study 1, there were significant memory changes under all conditions from the pre-scan test to the three-days-later test (*enhance dishonesty*: *t* = 4.87, *df* = 36, *p* <0.001***, 95% CI from 0.04 to 0.097; *enhance honesty*: *t* = 3.50, *df* = 36, *p <*0.0013**, 95% CI from 0.018 to 0.068; *random*: *t* = 4.45, *df* = 36, *p <*0.001***, 95% CI from 0.03 to 0.08; Table S6 in supplemental materials). We then investigated confidence and found that from pre-scan to post-scan test, significant confidence decline only occurred from the pre-scan to the three-days-later test (Figure 1D right panel; Table S10 in supplemental materials).

Moreover, the result of choice entropy was replicated in study 2 that there was a significant main effect of condition on choice entropy, with the *enhance honesty* condition lowest (*F*_3,108_ = 52.9, *p <* 0.001; 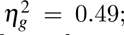 Figure S4C). The relationship between transformed confidence change and response **randomness** under *enhance dishonesty* and *random* conditions replicated the results in study 1 (*enhance dishonesty* & pre-post: β = -0.015, *p* = 0.20; *random* & pre-post: β = 0.0058, *p* = 0.30; *enhance dishonesty* & pre-three days later: *beta* = -0.01, *p* = 0.44; *random* & pre-three days later: β = 0.012, *p* = 0.30). We also plotted the dishonesty probability as the function of reward and the **trial-by-trial ‘consistency’**(see Figure S4D (right panel)), which confirmed the role of reward and consistency in dishonesty.

#### Replication of the multidimensional metacognition

Except for MAD, a significant decline of transformed variables from pre-scan to post-scan and three-days-later test were replicated in the *enhance dishonesty* condition (see Figure S5 in supplemental materials). To validate whether transformed RT and mouse tracking measurements reflected metacognition, we found that RT and mouse tracking measurements could predict transformed confidence in every stage of the memory test (Figure S9 right panel in supplemental materials). However, the change of transformed AD from pre-scan to three-days-later test couldn’t predict the transformed confidence change (Figure S9 right panel in supplemental materials). For PCA results, we found that in study 2, four components were needed to explain over 90% of the variance for the metacognition change from pre-scan to three-days-later test (Figure 4A), which was different from study 1.

#### Replication of univariate activations

The results of GLM analysis were replicated in the study 2. For GLM with response (dishonesty and honesty conditions) as the regressors, PrACC, vlPFC, DLPFC and retrosplenial cingulate cortex (RSC1) were also tracked under the dishonesty condition compared to the honesty condition (RSC1: β = -10.39 ± 18.49, *t* = -3.42, *p* = 0.0016; prACC: β = 3.80 ± 7.74, *t* = 2.99, *p* = 0.0051; vlPFC: β = 3.42 ± 6.91, *t* = 3.01, *p* = 0.0048; DLPFC: β = 3.19 ± 7.07, *t* = 2.74, *p* = 0.0094; Figure 2A lower panel). In the whole-brain map with a threshold of 0.01, significant clusters of DLPFC, preSMA, and insula were also found in dishonesty VS. honesty responses with *p <* 0.05 FDR corrected in supplemental materials), while PCC was found in honesty responses vs. dishonesty responses (Figure 2; see Table S12). Activations of two studies were plotted together, thus large brain areas of overlap were observed. For trials where the reward probability of incorrect choice was higher than correct choice, we didn’t observe a significant difference in reward-related ROIs like OFC either (β = 0.51 ± 11.53, *t* = 0.27, *p* = 0.79; Figure 2C upper panel) but in cognitive control related regions (MTG: β = 3.89, ± 7.21, *t* = 3.28, *p* = 0.0023; DLPFC: β = -2.53 ± 6.41, *t* = 2.40, *p* = 0.022; lTPJ: β = 8.91 ± 10.41, *t* = 5.20, *p <*0.001; Figure 2C lower panel).

#### Replication of separation effect of cumulative dishonesty in hippocampus

Except for OFC, the separation effect of consistent dishonesty was replicated in cognitive control brain areas like DLPFC and dACC, brain regions in the emotion system like lTPJ and insula, and memory systems like the hippocampus in study 2. Positive separation scores were observed in all ROIs (Figure 2D lower panel; for statistical details, see Table S14). Separation effects for dishonesty were also replicated in all ROIs except for PCC (Figure 2D lower panel; see Table S14 in supplementary materials). For reward, no separation effect was observed in any of the ROIs (Figure 2D lower panel; for other ROIs, see Table S14 in supplementary materials).

#### Replication of compressional rotated neural geometry of consistency and reward under dishonesty and dishonesty responses

The regression of model RDMs on neural RDM was partially replicated in study 2 that within all ROIs more than one model explained a significant amount of variance in the neural RDMs (see Table S16 in supplementary materials). In study 2, the neural geometries within ROIs were replicated. The compression along reward was significantly larger than consistency under dishonesty in all ROIs (SMA: *df* = 36, *t* = *−*7.36, *p <* 0.001; OFC: *df* = 36, *t* = *−*5.82, *p <* 0.001; lTPJ: *df* = 36, *t* = *−*6.10, *p <* 0.001; PCC: *df* = 36, *t* = *−*6.57, *p <* 0.001; insula: *df* = 36, *t* = *−*5.92, *p <* 0.001; Figure 3B upper right panel). For compression under honesty responses, we also replicated the results that consistency and reward didn’t have a significant difference (Figure 3B upper right panel; see Table S18 in supplementary materials for details). The estimated rotation parameter was not close to zero (SMA: *M* = *−*11.45; OFC: *M* = *−*11.07; lTPJ: *M* = *−*10.79; PCC: *M* = *−*12.14; insula: *M* = *−*8.05; Figure 3B right lower panel). And the third dimension encoded responses, indicated by a non-zero offset parameter, which were replicated in study 2 that offsets of all ROIs were around 1 (SMA: *M* = 0.995; OFC: *M* = 0.997; lTPJ: *M* = 0.996; PCC: *M* = 0.997; insula: *M* = 0.996; Figure 3B left lower panel). Taking OFC as an example, the final neural geometry was visualized by multidimensional scaling (MDS) (Figure 3C right panel), where the length of reward in spaced grids of dishonesty condition was shorter than consistency.

#### Replication of the relationship between the compression of reward under dishonesty responses and metacognition change

Correlations between compression ratio under dishonesty responses and immediate metacognition change were replicated in all ROIs (SMA: *rho* = 0.098, *p* = 0.0057; OFC: *rho* = 0.12, *p <* 0.001; lTPJ: *rho* = 0.063, *p* = 0.048; PCC: *rho* = 0.09, *p* = 0.012; insula: *rho* = 0.10, *p* = 0.0056; Figure 4B lower panel). From the pre-scan to three-days-later test, the relationship between metacognition change and compression ratio under dishonesty responses was also replicated that only in OFC the permutation test showed significant results (*rho* = 0.06, *p* = 0.05; Figure 4C lower panel).

#### Replication of functional connectivity reflecting the metacognition change

In study 2, the significant FCs between DLPFC and PCC, hippocampus were replicated (Figure 5A, right panel). FC between DLPFC and lTPJ could significantly predict immediate and delayed metacognition change as in study 1 (immediate: β = 1.70, *p <* 0.001, 95% CI from 1.09 to 2.31; delayed: β = 0.50, *p* = 0.03, 95% CI from 0.05 to 0.96; Figure 5B).

### Memory performance does not rely on visual or semantic features of stimuli in two studies

To examine whether memory change relies on visual or semantic features, we leveraged two DNNs, ResMem and VGG19. In contrast to a human participant, DNN predicts each image without being influenced by previous decisions or experimental settings. Thus, the resulting predictions are independent of local contextual effects introduced by the stimulus set. *Resmem* is a residual deep neural network (DNN) for predicting the intrinsic memorability of an image, which was trained on an independent image dataset^53^. A previous study has proven the validity of Resmem with that for predicting the memorability of another dataset in a group of participants that the predicted memorability score by Resmem is positively correlated with human memory performance^54^. Thus, we generated the memorability score for each food stimulus and tested whether there was a significant gap between conditions or correct and wrong stimulus. A two-way ANOVA was computed with conditions (*enhance dishonesty* / *enhance honesty* / *random*) and correctness (correct vs. incorrect) as independent variables. The results indicated that no significant main effect of condition and correctness on memorability score was found (condition: *F*(2, 66) = 1.18, *p* = 0.31, 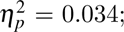 correctness: *F*(1, 66) = 0.72, *p* = 0.40, 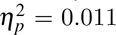). However, there was an interaction between the correctness and the condition (*F*(2, 66) = 3.37, *p* = 0.04, 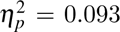). Post-hoc test showed that in the *enhance dishonesty* condition, the memorability scores of correct food pictures were higher than incorrect food pictures (*t* = 2.54, *p* = 0.019). For correct items, items of the *enhance honesty condition* were more memorable than those of the *enhance dishonesty* condition (*t* = 2.83, *p* = 0.02).

DNNs have shown effectiveness in predicting the neural responses of different regions in the visual system^55^. These studies highlight a crucial insight that earlier layers in the network represent low-level visual information such as edges, whereas later layers represent more complex and semantic features such as categorical information^56^. VGG19 is a variant of the VGG model which consists of 16 convolutional layers and 3 fully connected layers^57^. We aggregated 19 layers into 8 layers and extracted features of each layer to examine whether the feature representation of correct food items was different across conditions. Thus for each layer, we calculated the typicality score of each food item, which was defined as the similarity summation (Pearson correlation) of flattened features between the certain food item and all other food items (see *Methods* in supplemental materials)^54^. We compared typicality scores across three conditions at both early and late layers to check whether the correct items under a certain condition were visually or semantically prototypical to affect memory distortion. A two-way ANOVA was computed for each layer, taking condition and correctness as independent variables. Results showed that there was neither significant effect nor interaction of them (Table S4 in supplementary materials) on typicality score (Table S4 in supplementary materials).

We also examined whether the semantic similarity between correct food and error food differed across conditions via a well-established semantic model, Directional Skip-Gram^58^. The labels of foods were generated by five independent raters who did not participate in the experiment. This Chinese word2vector semantic model transformed the label of each food into a vector of semantic features, consisting of 200 values. The semantic similarity matrices between each pair of foods were obtained by calculating the cosine similarity of these word vectors. A one-way ANOVA was computed to examine the effect of conditions, and the results showed that there was no significant difference in semantic similarity among conditions (rater 1: *F*(2, 33) = 3.13, *p* = 0.057, 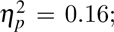 rater 2: *F*(2, 33) = 0.20, *p* = 0.820, 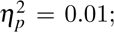 rater 3: *F*(2, 33) = 1.65, *p* = 0.207, 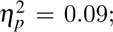 rater 4: *F*(2, 33) = 0.77, *p* = 0.471, 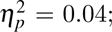 rater 5: *F*(2, 33) = 2.10, *p* = 0.139, 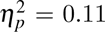).

## Discussion

Human memory is not simply a storage of our experiences or knowledge but also involves active awareness of remembering certain actions. Since evidence concerning how motivated repeated dishonesty affects metacognition is lacking, we were interested in exploring how the process of dishonesty influences metacognition changes. We investigate the relationship between neural responses of consistency and reward considerations in moral decisions and its link to metacognition change (Figure 6A). Our findings reveal a decline in memory retention and metacognition for immoral decision-related items, with associated neural representation differences in these decisions. The causes of this effect may be attributed to various factors, including the threat to self-image, the emergence of aversive emotions, or differences in the association of rewards. Here we show that the metacognition change is linked to the neural representation in the reward evaluation brain (OFC) during moral decisions (Figure 6C). It supports the fact that the post-decision effect on metacognition of memory depends on how reward is taken into account in moral decisions.

**Figure 6.**
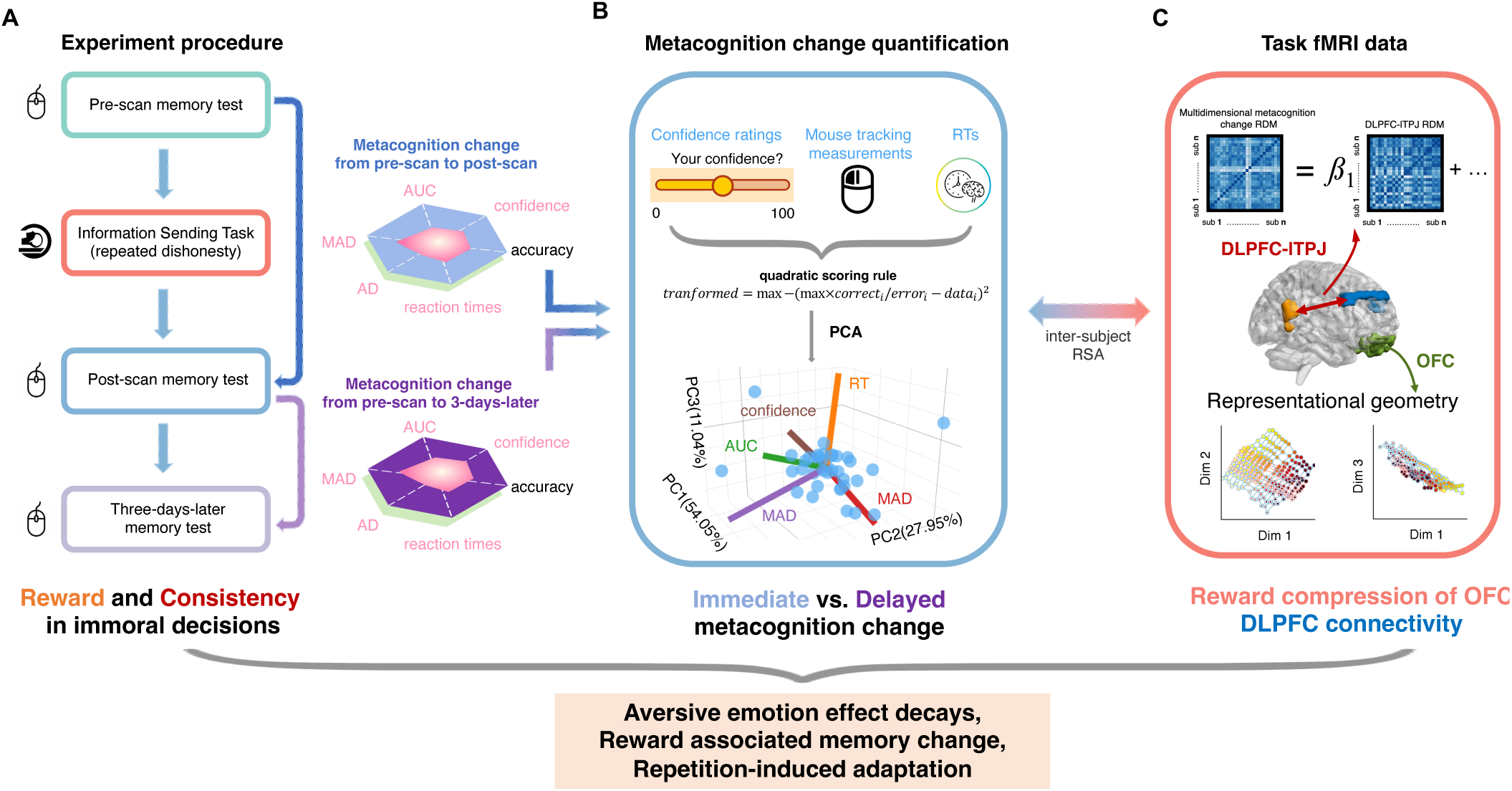
Summary: Neural mechanism of metacognition change induced by repeated dishonesty. We hypothesized that unethical amnesia might be due to the decay of aversive emotion, reward, and repetition-induced adaption. **(A)** We designed a multi-step experiment consisting of pre-scan and post-scan memory tests and information sending task, where participants were induced to lie. We recorded RTs, mouse tracking indices, and self-reported confidence in memory tests. **(B)** Metacognition change manifests on multiple dimensions of response in the pre- and post-scan memory tasks. These measurements all reflected metacognition levels after being transformed by the quadratic scoring rule. They jointly represent the immediate and delayed metacognition change after orthogonalization using PCA. **(C)** We found that unethical amnesia is the consequence of neural interactions between reward, emotion, and cognitive control. Metacognition change induced by repeated dishonesty is associated with the compression of reward in representational geometry in OFC. It supports the fact that the post-decision effect on metacognition of memory depends on how reward is taken into account in moral decisions. Functional connectivity (FC) between DLPFC and lTPJ during dishonesty can predict future metacognitive change. These results have implications for interpreting metacognition change after immoral decisions and the underlying cognitive processes that influence memory in social decisions.

### Metacognition change manifests in multiple responses in the memory task

The main finding of the current work is that memory and metacognition indeed change after dishonest behavior. Two pieces of evidence support the memory decline after immoral responses. First, we find that memory decline happens in the *enhance dishonesty* conditions (where participants lied more; see Figure S3B) characterized by a decrease in the accuracy and transformed confidence level in memory. Conversely, the effect was not found for the *enhance honesty* condition, indicating that the cause of amnesia can be attributed to unethical behavior. Previous studies investigating unethical amnesia mainly focused on memory error^59^ or self-reported memory^60^, but our results show that both accuracy and transformed confidence decreased in memory tests. We also combined other data dimensions, such as RTs and mouse tracking measurements, to quantify metacognition change comprehensively. From the dissonance perspective, this memory decline can be viewed as a manifestation of the impact of moral threats on one’s self-image^3, 61^. It may serve as an efficient strategy to address dissonance^19^. This perspective offers a compelling insight into the intricate interplay between moral integrity, self-perception, and memory, emphasizing the significance of moral considerations in shaping the metacognitive abilities of memory.

Notably, we validate the effectiveness of multidimensional variables in reflecting metacognition (Figure S9 in supplemental materials) and provide a more comprehensive quantification of metacognition change after dishonest responses (Figure 6B). Previous studies show indices of mouse tracking like MAD, AD, and AUC can be used as implicit manifestations of memory^62, 63^ as they reflect the conflict between two choices, thus suggesting the confidence level within participants. Also, the estimates of decision confidence in the metacognition task can rely on RT that longer RT suggests larger uncertainty^46^. However, in our preliminary results, we observe a decrease in RTs from the pre-scan to post-scan memory test (see Figure S3D in supplemental materials) reflecting a decrease in uncertainty, yet a decline in self-reported confidence rating (see Figure S4A in supplemental materials) and an increase in mouse measurements (see Figure S3 in supplemental materials). A possible explanation for this seemingly contradictory result is that people’s beliefs about their capabilities of memory performance(e.g., confidence) can be independent of the memory performance per se^64^. After mapping the statistical folded-X pattern of self-report confidence, RT and mouse tracking indices in a self-centered frame of reference using the quadratic scoring rule^9, 11, 12^, we observe a relatively replicable decline pattern of transformed variables in the *enhance dishonesty* condition. Also, the transformed RT and mouse tracking measures successfully predicted the transformed confidence (Figure S9 in supplemental materials), proving evidence that these indices can reflect metacognition levels.

### Immediate and delayed metacognition change after repeated dishonest and possible accounts

We find both immediate and delayed metacognition change effects, which can be explained from two aspects: emotion during the decision and repetition-induced adaptation. Confidence of memory changes may link to the emotion during decision, which has been shown to impact memory performance^65, 66^, and the decision-induced aversive emotion effect can be decayed over time^67^. Selfish lies harm moral self-image and people form self-serving beliefs to justify immoral acts before or after the decisions or actions^68^. There are several possible accounts for such unethical amnesia: i) the decrease in willingness to memorize due to the psychological discomforts^69^, ii)the confidence of the lied items would decrease as the inconsistent response^70^; iii) false memory as the repeated lies^71^ becomes truth with less conflict in the brain as the time lapses, which will not change the memory confidence of the lied items as the false memory elicits no conflicts.

Indeed, memories of harmful actions become obfuscated over time^72^, and we show that items in dishonest conditions decrease metacognition in different dimensions (Figure S5 in supplemental materials). Our observed metacognitive change can be related to the particular emotions induced by the decision context in which dishonest information can be passed in the fMRI scanner session. Indirect affective state measures and self-reports have not been taken. However, the behavioral results confirm that immoral individuals are more likely to adapt to dishonesty. From the replicable univariate results of dishonesty vs. honesty which exhibit a linear decline in cognitive control related brain regions (Figure S14 and Figure S15 supplemental materials), we can thus infer dishonest participants require less cognitive control process with less discomfort as the number of trials increases.

For the repetition-induced adaptation account, the participants can repeat instrumental behavior to achieve more rewards, even if the individual is not fully aware of the extent of the memory changes. First, we observe higher response randomness on *random* and *enhance dishonesty* conditions than on *enhance honesty* conditions (see Figure S4C). Compared to the *random* condition, repeated actions play a more significant role in decisions made in the *enhance dishonesty* condition (see Figure S3E in supplemental materials that the dishonesty rate is higher in the *enhance dishonesty* condition). However, there is no relationship between response randomness and transformed confidence change (see *Results* that response randomness did not predict transformed confidence change under all conditions). Thus, replicable accuracy and transformed confidence change effect (rather than self-reported confidence) occur in *enhance dishonesty* but not the *random* condition, partly ruling out the account that the memory change is due to the inconsistency of responses. This may contradict the prediction that repetition would not affect metacognition judgment strongly^73^, and previous evidence shows repetition can increase confidence in judgment^74^. As for our findings of behavioral variables combining brain evidence that could reflect conflict level presented a contradiction, we can only conclude that false memory as the repeated lies becomes truth via decreased metacognition. There is still a lack of evidence explaining its causes, especially regarding confidence.

### Reward compression of OFC in neural geometry correlates with immediate or delayed metacognition change

As for the role of reward in immoral decisions, our fMRI results imply a compression of reward (compared to consistency), especially for dishonesty condition, over emotion and memory ROIs (e.g., SMA, OFC, lTPJ, PCC, and insula). This finding is also confirmed by behavior results in DDM, which suggest a smaller drift rate than consistency (see Figure S13). Moreover, we find that reward compression of OFC in neural geometry is correlated with both immediate and delayed metacognition change, and this result is replicated in Study 2 (Figure 4B and C). The involvement of OFC in both immediate and delayed metacognition change suggests that this brain region may be crucial in mediating the impact of reward-related neural activity on distortions in metacognition over time. This finding echoes the previous neurophysiological work in primates that OFC plays an important role in subjective value encoding^75, 76^, reward processing, memory-guided decision making^77, 78^. It is also consistent with the evidence indicating OFC is one candidate neural substrate for propositional confidence^79–81^. Although considerable evidence indicates that the PFC (including orbital PFC) and hippocampus show bidirectional flow of information^82^, we have not replicated the results for the hippocampus associated with immediate and delayed metacognition change, with evidence in study 1 that reward compression of the hippocampus in neural geometry was correlated with both immediate and delayed metacognition change, but not study 2 (Figure S12 in supplemental materials). Other possible accounts exist, and we can not exclude the other ways to mediate such metacognition change, such as the interplay among memory, reward, and cognitive control in the task that may lead to metacognition change.

### DLPFC connectivity in moral decision-making predicts metacognition changes

Our results reveal a robust and replicable functional connectivity between DLPFC and lTPJ in predicting immediate and delayed metacognition changes (see Figure 5B) after repeated dishonesty. These results suggest that the interaction between DLPFC, a key component in cognition control and dishonesty^50^, and lTPJ which involves guilt processing^83, 84^ and self-image processing^50^, plays a crucial role in predicting the extent of metacognition change following dishonest behavior. Consistent with previous studies, moral versus immoral decisions are driven by both rewards and self-image^33, 85–87^. Dishonesty involves cognitive-control related brain regions like DLPFC and anterior cingulate cortex (ACC). DLPFC plays a key role in dishonesty and cognitive control, and the brain lesion^88–90^ in DLPFC can increase dishonesty, while stimulation over DLPFC can decrease it^91^. TPJ, involved in self-processing network, is also responsible for maintaining a positive self-image and thereby increasing honesty^50^. Harmful actions elicited a higher response than neutral acts in lTPJ^92^, and restraining the rTPJ or lTPJ with tDCS decreased the role of beliefs in moral judgment^93^.

The DLPFC and lTPJ are two brain regions closely related to metacognition. Previous studies revealed the role of DLPFC in mediating confidence judgments in visual memory^94^, and DLPFC was especially recruited in re-decision phase^95^. lTPJ has been implicated in the Theory of Mind reasoning^96–98^, and lTPJ might play a vital role in mediating self-other mergence during mentalizing^99^. Moreover, evidence from studies in anosognosia for hemiplegia (AHP) has revealed that the disorder of cognitive systems involved in prospective belief updating (prospective metacognition) or self-referentiality was associated with significant disconnection of TPJ, insula and prefrontal cortex^100–102^. These associative cortical regions were connected through several white matter tracts, connecting regions of different cognitive networks like default, ventral attention, and cingulo-opercular^101^. This echoed our results that connectivity between DLPFC and lTPJ during dishonesty task could predict later metacognition change, suggesting that metacognition change induced by repeated dishonesty extended beyond local, retrospective monitoring. Moreover, brain lesion studies identify distinct network connections associated with visual and motor anosognosia and a shared, cross-modal network for awareness of deficits centered on memory-related brain structures^103^. Together, metacognition relies on a distributed network rather than certain brain regions, emphasizing the pivotal role of neural interactions between different neural systems in shaping self-awareness.

In summary, our findings imply that the neural mechanisms governing both reward processing (involving the OFC), cognitive control (involving DLPFC) and self-processing network (involving lTPJ) are intricately involved in shaping the long-term effect of dishonest behavior. This finding sheds light on the neurological underpinnings of metacognition change in the context of moral decision-making, emphasizing the complex interplay between neural activity and ethical behavior. Thus, it provides valuable insights into the cognitive processes involved in moral decisions and their interplay with memory awareness. It is also interesting to identify the specific dynamic mechanism of such memory change effect, with potential implications for addressing issues related to unethical behavior and self-integrity.

### Limitations, future work, and conclusion

Notably, existing studies have interpreted unethical amnesia as a result of self-image keeping, inconsistent responses induced metacognition change, or due to evidence of conflict and uncertainty. We find that metacognitive change is primarily related to reward compression under dishonesty, and the extent to which participants’ representational pattern of the decision information. However, based on these findings, we still can not exclusively interpret unethical amnesia as a result of representational changes of decision information representation in dishonesty as other accounts may contribute to it. For example, we use food preference in the task and find out the tendency towards the correct answer. Thus, the conclusion may be tied to a specific domain. This finding suggests that future work can develop different domains of decision tasks covering choices from preference, to other types of autobiographical information. Further, most previous studies have measured confidence in memory before completing a memory task, and measuring it after the task may increase their capacity or decrease the measure validity. Future research could test our predictions with various confidence measures and confirm the validation of our results.

In summary, we reveal a vital neural response linked to metacognition change in memory after immoral decisions with reward-seeking and self-image considerations. We find repeated immoral decisions get faster, but are still linked to the cognitive control brain. The metacognition change to unethical responses is associated with representational geometry of the compression ratio between consistency and reward in OFC, but not with the visual or semantic item features. These results have implications for interpreting metacognition change after immoral decisions and the underlying cognitive processes that influence memory in social decisions.

## Methods

### Lead contact

Further information and requests for resources and reagents should be directed to and will be fulfilled by the Lead Contact, Haiyan Wu

### Data and code availability

Behavioral data and accompanying code for all behavioral analyses and figures can be found on the Open Science Framework (https://osf.io/c4qbr) or GitHub (https://github.com/andlab-um/Unethical-Amnesia).

### Materials availability

Materials are available from the corresponding author upon reasonable request.

### Participants

26 healthy participants (16 females, 10 males, mean age = 20.12 ± 2.47 years old) took part in study 1. For the replication study (study 2), 37 healthy participants (31 females, 6 males, mean age = 23.27 ± 3.52 years old) were recruited in Macau. Participants were recruited from the student community at the University. All participants were right-handed and had no history of neurological or psychiatric disorders. Participants gave written informed consent after a complete description of the study was provided. Participants received a full debriefing on the completion of the experiment. All the procedures involved were by the Declaration of Helsinki and were approved by the Institutional Review Board (IRB) of Caltech (IRB 18-0829) (study 1) and the University of Macau (BSERE21-APP005-ICI). The sample size for the fMRI study was determined based on the previous fMRI studies on dishonesty (e.g., n = 28^21^, n = 40^50^, n = 25^104^, n = 31^37^). In addition, we recruited five raters who did not participate in the experiment to provide labels for food pictures used in this study.

### Stimuli

We implemented two sets of information (Personal information vs. Food preference) about a stranger for all participants. For additional methodological details, please refer to the supplemental materials.

#### Personal information

We created a set of personal information for two people with which there is a true answer for one, but a false answer for another person. Table S1 (see supplemental materials) provides the personal information and both of the information are foreigners for the participants of the current study, to control the familiarity or social effects.

#### Food preference

We selected a set of 72 food images to use as the memorization materials from the FoodPic database^105, 106^. Some examples of foods are listed in Figure S1. 36 food images were printed as the preferred foods of Mr. Li (a common last name in Chinese) to memorize (food image subset 1) while another 36 food images were used as the irrelevant food or non-preferred food. The usage of two subsets is randomly assigned to the participant to control the effect of the materials in the task. The labels of the pictures, which were further used in Directional Skip-Gram, were generated by five independent raters who did not participate in the experiment.

### Study design and procedure

The whole study is a within-subject design, with 9 steps (8 steps on the fMRI scanning day and 1 step for 3 days after the MRI scanning day) in total. The information task was inspired by previous studies that examined self-serving dishonesty involving information passing and advice giving^21, 107, 108^.

After arriving at the lab for the fMRI task, all participants were told that there are several steps in information receiving and passing in the experiment. They read the instructions for each step and started the specific task after understanding all of the information (the details of the instructions are provided in the supplemental materials).

Participants were told that the information to receive or memorize was randomly assigned by drawing an envelope from two envelopes numbered 1 and 2, and that all participants were going to do it before the experiment. They were told that the experimenter would write down the stimuli set number and they should not open the envelope until step 3. In reality, participants always received the food image materials which will be further used in step 3.Except from the information sending task (IST; see below), all mouse trajectories of the binary response were recorded.

#### Step 1: The first food preference task with mouse tracking

In the first step, participants were asked to make a binary choice to choose the food they preferred between two choices, and then rate the preference level (Figure 1A). They were told that there weren’t any consequences for their choices in the preference task. A mouse tracking approach was implemented in this task to record their mouse trajectories, helping the subjects adapt to the mouse tracking approach which would be repeatedly used in this study.

#### Step 2: Information (personal information of a stranger) receiving and trust decisions with mouse tracking

In this step, participants were asked to check the personal information (e.g., name, best friend name, name of the pet, etc.) received about a foreigner, whose information was sent from a previous anonymous participant. The information sender memorized the personal information and sent it by mouse clicking, while the participant did not know the person and has zero information about the true personal information. The received information was received with two alternative items, one of which is with an asterisk indicating that the information sender (who has memorized the information) clicked it and sent it to the participant. The participants were asked to click the correct answer to gain more earnings, even though they were given limited information except the marked answer from the sender. There were 24 trials in total, which lasted around 5 minutes. To exclude the emotional effect on the following steps, note that no feedback would be delivered until the end of the whole study. There are three aims to designing this step: 1) to make the participant believe in the protocol of information passing, in which they will receive information from others and send memorized information to other participants as well; 2) by using this 0-information context, we can measure the baseline trust level, that how much a participant trust others by clicking the indicated items (marked as asterisk) in the task.

#### Step 3: The first food preference memory task with mouse tracking

During this step, all participants were asked to open the envelope and memorize Mr. Li’s preferred foods with 90% accuracy. For this sufficient memorization task, one needs to engage in extensive practice to achieve nearly perfect accuracy about another one’s food preferences. We measured their binary response within the paired foods, and the confidence of the memory (see Figure 1A). To get a baseline of correct memory hand trajectory for each item, participants were asked to do the memory test with mouse tracking and click the correct answer for the memorized preference quickly. There was no feedback in each trial, but the total accuracy would be delivered at the end of the test. After reaching an accuracy of more than 90% (all participants can reach this in the test after practice), participants can continue to the next step.

#### Step 4: The information sending task (IST) in the fMRI scanner

In the fMRI task, participants were asked to pass the preference information of Mr. Li to the next participant (the receiver) with the consideration of reward units. The receiver may get more reward units if they click more correct answers based on the sent information. When considering the higher reward corresponding to the incorrect answer, participants may pass the incorrect answer to the next participant (dishonesty). We manipulated reward unit difference (referred to as “reinforcer”in this paper) between two items and kept them lying about specific items (repeated dishonesty) or making truthful responses with 4 sessions of fMRI scanning.

For the participant, the motivation to make a dishonest response is to gain more reward units for themselves, which is self-serving. To manipulate the level of consistency **(quantified as the difference of cumulative history responses between incorrect choice and correct choice)** and response **randomness (choice entropy)**, we set three conditions with 12 food items each. The first condition, *enhance dishonesty*, is to induce consistent dishonest responses by consistently keeping higher reward units for incorrect answers. The second condition, *enhance honesty*, is to reinforce honest responses, by keeping a higher reward unit for the true answer while with a lower reward unit for the incorrect answer. The third condition is the *random* condition, with random reward units’ difference of two answers, which means the participant may give random responses upon the rewards or keep consistent responses or honest responses across sessions. Thus three conditions contain consistent honesty, consistent dishonesty, and the level in between.

The reward unit differences were drawn from a uniform distribution (2:2:8 units) with pseudo-random sampling with replacement, which is well-matched for all three conditions. For each condition, we repeated 8 times for each item, which with 2 times for each scan session, thus participants played a total of 4 sessions with 72 trials each (36 pairs of food images in each round and 2 rounds in each session) for a total of 288 trials. After presenting both the food item and the reward unit corresponding to the answer on the left and right sides, the participant was asked to decide to send information within a duration of 4s time limit. The sent item was highlighted for 1s, whereas jitters between trials were 2–8 s (M = 4 s) long. Further, it has to be noted that although our design informs the participants that their responses will be sent to other participants, the responses with either correct or incorrect answers in repeated 8 trials might not seem very realistic as they only received other’s information once in step 2. At that time, they did not know they would conduct step 6 (see below) later on.

#### Step 5: Exit questionnaire

To acquire an explicit measure of the motives and make the natural “forgetting”by making participants believe that the experiment is almost finished, we have an exit questionnaire and a “faked”debrief part. That is, we tell the participant they almost finish most of the task, and then go to finish the questionnaire. In this step, participants will finish questionnaires, such as interactive mentalizing questionnaire (IMQ), moral foundation questionnaire, psychopathy trait, autism trait, anxiety, and some debrief questions, like how much they believe in others, how much they think the other’s will trust their message, and their feelings and the aim of the study. This step lasted around 30 minutes.

#### Step 6: the surprise second memory test with mouse tracking

After step 5, participants were asked to do the memory test surprisingly. In this task, they will be asked to always click the correct answer for the memorized foods, and their confidence about the memory. More correct answers mean more bonus. There was no feedback until the end of the test.

#### Step 7: The second receiving and trust decisions with mouse tracking

This step was used to probe the possible trust change effect after the honest/dishonest responses in the IST or with more direct experience of how individuals send this information as they performed in the scanner. The procedure and instruction were the same as step s, participants were asked to click the correct answer to gain more earnings, with the marked answer from the anonymous sender.

#### Step 8: The second food preference task with mouse tracking

This step was the same as step 1, where participants were asked to make a binary choice to choose the food they preferred between two choices, and then rate the preference level. They were told that they didn’t need to consider what they had chosen in step 1 and there weren’t any consequences for their choices. Then, participants were informed that there would be another task three days later, however, note that they were not informed that there would be a memory test.

#### Step 9: Three days after the scanning: the third memory test with mouse tracking

Three days after the scanning time, participants were asked to do the memory test again. This step was the same as step 3 and step 6. In this task, we asked participants to click the correct answer for the memorized preferences of Mr. Li, and their confidence about the memory. More correct responses mean more bonuses. There was no feedback in each trial until the end of the test.

After all of the steps, participants will be paid with a list of performances in each step, in which their reward units are related to their real choices in step 4 and memory performance. The average payment for each participant is 150 yuan(ranging from 124-160 yuan).

### Manipulation check

We performed manipulation checks on the core manipulations in this study. In the post-scanner task, we asked participants whether they perceived another information receiver as real or not, and to guess the goal of the current study, the effectiveness of the experiment manipulation, or suggestions. On the perceived reality of another partner (1: yes, 2: no), all participants (26 out of 26) in study 1 believed the other player was real to them. In study 2, 25 out of 37 participants believed the other player was real to them (the level of trust was above 67%). None of the participants accurately guessed the aim of the study, see some example answers in the supplemental materials.

### Metacognition measurements

#### Mouse Trajectory measurements

We extracted mouse trajectories from pre-scan memory task and post-scan memory task for each participant. Standard mouse-tracking preprocessing was conducted temporally and spatially^17, 62^. In a typical binary choice design, trajectories end at either the left or the right response option. As the overall spatial direction is irrelevant for most analyses, all trajectories were remapped so that they would end on the same side. We used the R package ‘mousetrap’^109^ to map the trajectories to the left by default, suggesting that trajectories that end on the right-hand side are flipped from right to left. We rescaled all mouse trajectories into a standard coordinate space (top left: [-1, 1]; top right: [1,1]) so that the cursor always started at [0,0]^17, 110^. Temporally, time normalization was applied to the trajectories such that the duration of each trial was divided into 101 identical time bins using linear interpolation to obtain the average of their length across multiple trials^110–115^.

Several different measures including MAD, AD, and AUC for the curvature of mouse trajectories that have been proposed in the literature^62, 63^ were calculated. Specifically, AUC represents the geometric area from the reference trajectory to the actual trajectory. The higher the AUC is, the bigger the attraction to the other response option^62^. AD is sensitive to the temporal dynamics of the movement^116^. Such metrics offered a fine-grained quantification of the conflict or uncertainty among response options^117^.

#### Metacognition change calculation

According to the folded-X pattern, maximal metacognition is obtained both when one is maximally confident and right and when one is minimally confident and wrong. Five indices including confidence, RTs, and mouse trajectory measurements (MAD, AD and AUC) of pre-scan, post-scan and three-days-later memory tasks were transformed using the quadratic scoring rule (QSR)^11, 12, 47, 48^ to measure metacognition together.

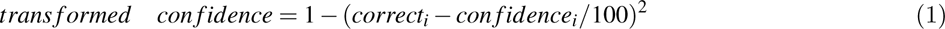

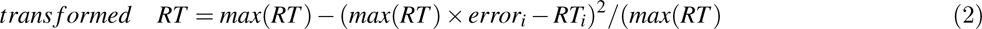

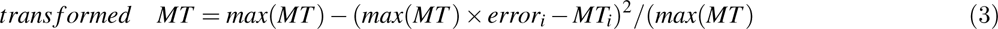

where correct_i is equal to 1 in a trial if the choice was correct and 0 otherwise, and error_i is equal to 1 in a trial if the choice was incorrect and 0 otherwise. Also, confidence_i, RT_i, and MT_i are the subject’s confidence ratings, RT, and mouse trajectory measurements (AUC/MAD/AD) on trial i. The quadratic scoring rule jointly considered accuracy and confidence, describing how closely participants’ confidence tracks their performances^11, 12, 48, 118^.

Now changes from pre-scan to post-scan and to three-days-later test of five transformed indices were averaged among trials per subject and were used to measure metacognition change together. Due to the co-linearity between mouse trajectory measurements, and the different scales of these five transformed indices, we conducted principal component analysis (PCA) on these indices and selected the components required to explain 90% of variance to evaluate metacognition change.

### MRI data acquisition

For study 1, MRI Data from four sessions were collected with a General Electric 3T scanner (GE Discovery MR750) in Beijing. Neuroimaging data acquisition included the collection of both anatomical and functional data. High-resolution T1-weighted structural image data were collected for anatomical reference using a 3D magnetization-prepared rapid gradient-echo (MPRAGE) sequence (voxel size = 1 mm isotropic; FOV = 192 x 256 mm; slice thickness = 1 mm; TR = 6.65 ms, TE = 2.93 ms, TI = 450 ms, flip angle = 12°; in-plane matrix resolution = 192 × 256). Whole-brain functional imaging was acquired by a T2*-weighted gradient echo, echo-planar pulse sequence in ascending interleaved order with a 4.2 mm slice gap, voxel size of 3.5 x 3.5 x 4.2 mm^3^, slice thickness of 3.5 mm, TE of 30 ms, TR of 2000 ms and flip angle of 90°.

For the replication study (study 2), MRI data were acquired using a 3.0 T Siemens MAGNETOM Prisma MRI scanner with a 64-channel head coil in the Center for Cognitive and Brain Sciences, University of Macau. Neuroimaging data acquisition included the collection of both anatomical and functional data. High-resolution T1-weighted images were acquired for each participant (3D MPRAGE sequence; voxel size = 1 mm isotropic; FOV = 176 x 256 mm; slice thickness: 1.0 mm; TR = 2300 ms, TE = 2.26 ms, TI = 900 ms, flip angle = 8°; in-plane matrix resolution = 176 × 256). Whole-brain functional imaging was acquired by a T2*-weighted gradient echo, echo-planar pulse sequence in ascending interleaved order with a 2.0 mm slice gap, voxel size of 2.0 mm, slice thickness of 2.0 mm, TE of 30 ms, TR of 1000 ms and flip angle of 90°.

### fMRI GLM analysis

GLM analysis was conducted via nilearn^119^. We performed three GLMs with four scan sessions (8 blocks) together. The time foods showed up on the screen was treated as the onset time.

To investigate the effect of dishonesty compared to honesty, our first GLM (termed GLM-1) specified moral responses (dishonesty versus honesty) as the regressor. The second GLM (termed GLM-2) split trials into two types in consideration of whether participants chose the option with a higher probability of getting the reward, validating the reward manipulation. Before conducting the GLM-1 and GLM-2, the images were smoothed using a 6-mm (study 1) / 4-mm (study 2) full width at half maximum (FWHM) Gaussian kernel. After that, they were masked using the anatomical mask generated by fMRIPrep for each session per participant.

The third GLM (termed GLM-3) was constructed for representational geometry analysis with unsmoothed preprocessed images. Conditions of GLM-3 were for each combination of moral decisions (dishonesty versus honesty), consistency difference (-7 to 7; might be different for each participant), and reward difference (-8, -6, -4, -2, 2, 4, 6, 8). The resulting GLM was convolved with SPM’s canonical hemodynamic response function. The model was corrected for temporal autocorrelations using a first-order autoregressive model and a standard high-pass filter (cutoff at 128 s) was used to exclude low-frequency drifts. To account for residual variance, 18 (estimated headmotion parameters, global signal, framewise displacement, six anatomical CompCor confounds and four discrete cosine-basis regressors) of the confounds generated by fMRIprep was selected to add to GLMs and a constant regressor for each entire session.

### Separation score

The separation scores were calculated based on GLM-3 to reflect the effect of responses (dishonesty VS. honesty), consistency (the number of history responses for the incorrect answer > the number of history responses for the correct answer VS. the number of history responses for the incorrect answer < the number of history response for the correct answer) and reward (reward probability for the incorrect answer > reward probability for the correct answer VS. reward probability for the incorrect answer < reward probability for the correct answer). It equals to the average dissimilarity in activity elicited within the condition compared to the average dissimilarity across conditions^120–122^. Hence, a significantly positive separation score for any of the factors (dishonesty, consistency, and reward) indicates that the ROI encodes representations of that factor.

### ROI selection

Cognitive control plays a key role in (dis)honesty to override the moral default^50, 52^ and keep self-consistency^33^. Our main task in fMRI scanner employed rewards to motivate individuals to lie, so we first selected ROIs involving cognitive control and reward processing. DLPFC was closely related to cognitive control, and the brain lesion^88–90^ in DLPFC can increase dishonesty, while stimulation over DLPFC can decrease it^91^. Another important brain region involved in moral decision was insula. Neuroimaging studies have shown that the insula is involved in moral judgment^123^, and detecting norm violations^124, 125^. For reward processing, a meta-analysis^104, 126^ investigated 142 neuroimaging studies and showed that mOFC, and PCC all respond more strongly to positive than negative monetary outcomes. Also, previous neurophysiological work in primates that OFC plays an important role in subjective value encoding^75, 76^

Apart from reward processing, previous literature on moral decision-making illustrates that moral versus immoral decisions are driven by both rewards and self-image^33^. Previous task-based fMRI studies suggest that the self-processing network consisted of mPFC, PCC, and TPJ, and might maintain a positive self-image and thereby increase honesty^50^. For dishonesty-induced memory change, we selected the hippocampus for its pivotal role in memory^127^. For metacognition change, we selected dACC because dACC encoded generalizable metacognitive control signals in various kinds of tasks^46, 128^.

### Representational similarity analysis with parameterized model

To better depict the neural geometry, we fit the neural RDM to a parameterized model where the neural geometry could be among the states of the 12 models described above. In the parameterized model, there were six free parameters: four controlling compression of consistency and reward under different responses, one controlling the angle between two grids, and one controlling the separation of two grids. Thus the full model was:

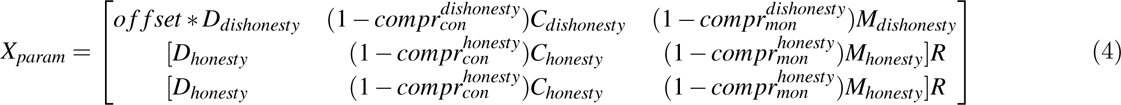

We fit the parameterized model to neural RDMs using a constrained optimization procedure (fmincon in MATLAB) with a least-squares cost function. As the procedure is sensitive to the choice of starting values, we averaged over 1000 independent runs with random starting values. The fitted parameters were used to visualize the representational geometries of the best-fitting RDMs via projection into three dimensions with classical Multi-Dimensional Scaling (MDS). The compression ratio was defined as the compression of consistency divided by the compression of reward.

### Permutation test

To explore the relationship between memory distortion and the compression ratio under dishonesty responses, inter-subject pairwise Euclidean distance was calculated by *Scipy*’s Spatial module to carry out a dissimilarity matrix for both memory distortion and compression ratio under dishonesty responses. Then we computed Spearman’s rank-order correlations between the condensed memory distortion dissimilarity matrix and the condensed compression ratio dissimilarity matrix. To obtain significance levels of the resulting Spearman’s *rhos*, we performed the permutation tests by shuffling the order of the observations in the memory distortion dissimilarity matrix 10,000 times, and calculated the proportion of the permuted *rhos* that exceeded the true *rho*.

### Functional connectivity analysis

To explore how the value signal in the BA9-DLPFC interacts with other ROIs during dishonest responses in the last session, we implemented a functional connectivity analysis using the CONN toolbox^129^ (version 21.a) (https://www.nitrc.org/projects/conn).

We employed a standard pipeline by using the pre-processed fMRI data. In the denoising step, we used linear regression to remove the influence of the following confounding effects on the fMRI time course: (1) BOLD signal from the white matter and CSF voxels (five components each, derived using the anatomical component-based correction (aCompCor) implemented using the ART toolbox), (2) eighteen confounds by fMRIPrep: six residual head motion parameters, global signal (the average signal within the brain mask), framewise displacement (the quantification of the estimated bulk-head motion calculated using formula proposed by^130^), six additional noise components calculated using anatomical CompCor and four DCT-basis regressors, (3) effect of task-condition using separate run regressors (for dishonesty and honesty conditions) convolved with the hemodynamic response function.

Thus, we performed the connectivity analysis on the residuals of the BOLD time series after removing condition-related activation/deactivation effects^129, 131, 132^. Finally, the denoising step included temporal bandpass filtering (0.008–0.09 Hz), and linear detrending of the functional time course. Following pre-processing, we performed condition-dependent functional connectivity analysis on the mean BOLD time course^129^ extracted from selected ROIs.

To investigate how functional connectivity calculated above could predict memory distortion, we constructed inter-subject representational dissimilarity matrix (RDM) of memory distortion (using the first four components) and functional connectivity (DLPFC-SMA, DLPFC-lTPJ, DLPFC-PCC, DLPFC-OFC and DLPFC-hip) using Euclidean distance^133, 134^. We regressed memory distortion over the rest four FC RDMs by using *lm* function in R.

### Data and code availability

FMRI data and analysis code are available from the corresponding author on reasonable request.

## Author contributions

HW contributed to the conception and design of the experiment, HW and XJX conducted the experiment and collected the data, XJX and HW analyzed the data, and XJX, HW, and DM drafted the manuscript. All authors reviewed the manuscript and gave final approval for publication.

## Acknowledgements

This work was mainly supported by the Science and Technology Development Fund (FDCT) of Macau [0127/2020/A3, 0041/2022/A], the Natural Science Foundation of Guangdong Province (2021A1515012509), Shenzhen-Hong Kong-Macao Science and Technology Innovation Project (Category C) (SGDX2020110309280100), MYRG of University of Macau (MYRG2022-00188-ICI) and the SRG of University of Macau (SRG2020-00027-ICI). We thank Quanying Liu, Xiaoqing Hu, and Wanjun Lin for their early comments on the manuscript. We thank He Miao, Shuhan Zhang, Hanran Luna Li, Haofei Wu, Yizhe Chen, and Yan Tian for their help in data collection. We also thank all participants who took part in the study and enabled this research to be possible.

